# Multi-omics Quality Assessment in Personalized Medicine through EATRIS

**DOI:** 10.1101/2023.10.25.563912

**Authors:** EATRIS Plus Multi-omics working group and stakeholders (in alphabetical order by last name), Patricia Alonso-Andrés, Davide Baldazzi, Qiaochu Chen, Elisa Conde Moreno, Lorena Crespo-Toro, Kati Donner, Petr Džubák, Sara Ekberg, Maria Laura Garcia-Bermejo, Daniela Gasparotto, Bishwa Ghimire, Jolein Gloerich, Janine Habier, Marián Hajdúch, Rashi Halder, Sari Hannula, Hanna Lindgren, Yaqing Liu, Roberta Maestro, Tom Martin, Pirkko Mattila, Lukáš Najdekr, Kenneth Nazir, Anna Niehues, Anni I Nieminen, Jessica Nordlund, Emanuela Oldoni, Elin Övernäs, Aino Palva, Maija Puhka, Ileana Quintero, Miren Edurne Ramos-Muñoz, Esperanza Macarena Rodríguez-Serrano, Sabrina Saracino, Andreas Scherer, Leming Shi, Jarmila Stanková, Peter-Bram ’t Hoen, Tanushree Tunstall, Beatrice Valenti, Alain van Gool, Marjan Weiss, Bhagwan Yadav, Yuanting Zheng, Patricia Žižkovičová

**Affiliations:** Biomarkers & Therapeutic Targets Group, Ramon and Cajal Health Research Institute (IRYCIS), Madrid, Spain; Centro di Riferimento Oncologico di Aviano (CRO Aviano) IRCCS, Via Gallini 2, 33081 Aviano (PN), Italy; State Key Laboratory of Genetic Engineering, School of Life Sciences, Human Phenome Institute and Shanghai Cancer Center, Fudan University, Shanghai, China; Institute for Molecular Medicine Finland FIMM, Tukholmankatu 8, Helsinki, Finland; Institute of Molecular and Translational Medicine, Faculty of Medicine and Dentistry Palacky University and University Hospital Olomouc, Hněvotínská 1333/5, 779 00 Olomouc, Czech Republic; Department of Medical Sciences and Science for Life Laboratory, Uppsala University, Box 1432, BMC 751 44 Uppsala, Sweden; Radboud University Medical Center, Department of Genetics, Translational Metabolic Laboratory, Geert Grooteplein zuid 10, 6525GA Nijmegen, The Netherlands; Luxembourg Center for Systems Biomedicine, University of Luxembourg, 7, Avenue des Hauts-Fournaux, Esch-sur-Alzette, Luxembourg; EATRIS ERIC, European Infrastructure for Translational Medicine, 1081 HZ Amsterdam, The Netherlands; Luxembourg Institute of Health, 1A-B, rue Thomas Edison, L-1445 Strassen Luxembourg; Radboud University Medical Center, Department of Biomedical Sciences, Geert Grooteplein zuid 10, 6525GA Nijmegen, The Netherlands

**Keywords:** multi-omics, quality, EATRIS, MOTBX

## Abstract

Molecular characterization of a biological sample, e.g., with omics approaches, is fundamental for the development and implementation of personalized and precision medicine approaches. In this context, quality assessment is one of the most critical aspects. Accurate performance and interpretation of omics techniques is based on consensus, harmonization, and standardization of protocols, procedures, data analysis and reference values and materials. EATRIS, the European Infrastructure for Translational Medicine (www.EATRIS.eu), brings together resources and services to support researchers in developing their biomedical discoveries into novel translational tools and interventions for better health outcomes. Here we describe activities of member facilities of EATRIS towards quality assessment of pre-clinical sample processing, clinical omics data generation, multi-omics data integration, and dissemination of the resources in a Multi-Omics Toolbox, the principal deliverable of the EATRIS Plus project for the consolidation of EATRIS towards translational Medicine.

## Introduction

Precision medicine relies on sensitive and specific detection of biological variables that may support diagnosis, prognosis, and prediction of therapy response. In this context, omics technologies, which detect and quantify the abundance of thousands of molecular entities such as nucleic acids, proteins, and metabolites in a given biological sample, have proven invaluable tools. Moreover, two or more omics technologies may be integrated into multi-omics approaches, thus increasing the possibility of detection of molecular signatures, and providing a more comprehensive view of the molecular portrait of the specimen. Clearly, the accuracy of such type of molecular diagnosis demands a high quality of data. This is becoming increasingly crucial, given the widespread use on omics approaches in routine diagnostics and the progressive expansion in the clinical applications. In addition to making data findable, accessible, interoperable, and reusable (FAIR-principles, https://www.go-fair.org/fair-principles) (1, 2), assuring a minimal level of data quality is essential for further use towards the benefit of individuals in healthcare and clinical care.

Integrated multi-omics data analysis and interpretation require careful design of experiments and associated data analysis procedures to enable optimal use of research resources. Importantly, quality needs to be assessed and maintained throughout the whole process of generation and analysis of multi-omics approaches. In this regard, several initiatives have been undertaken to promote best practices, ranging from a proper definition of the experimental question and study design, sample handling, data analysis and stewardship, and re-use of approaches and data (**Figure 1)**.

**Figure 1.**
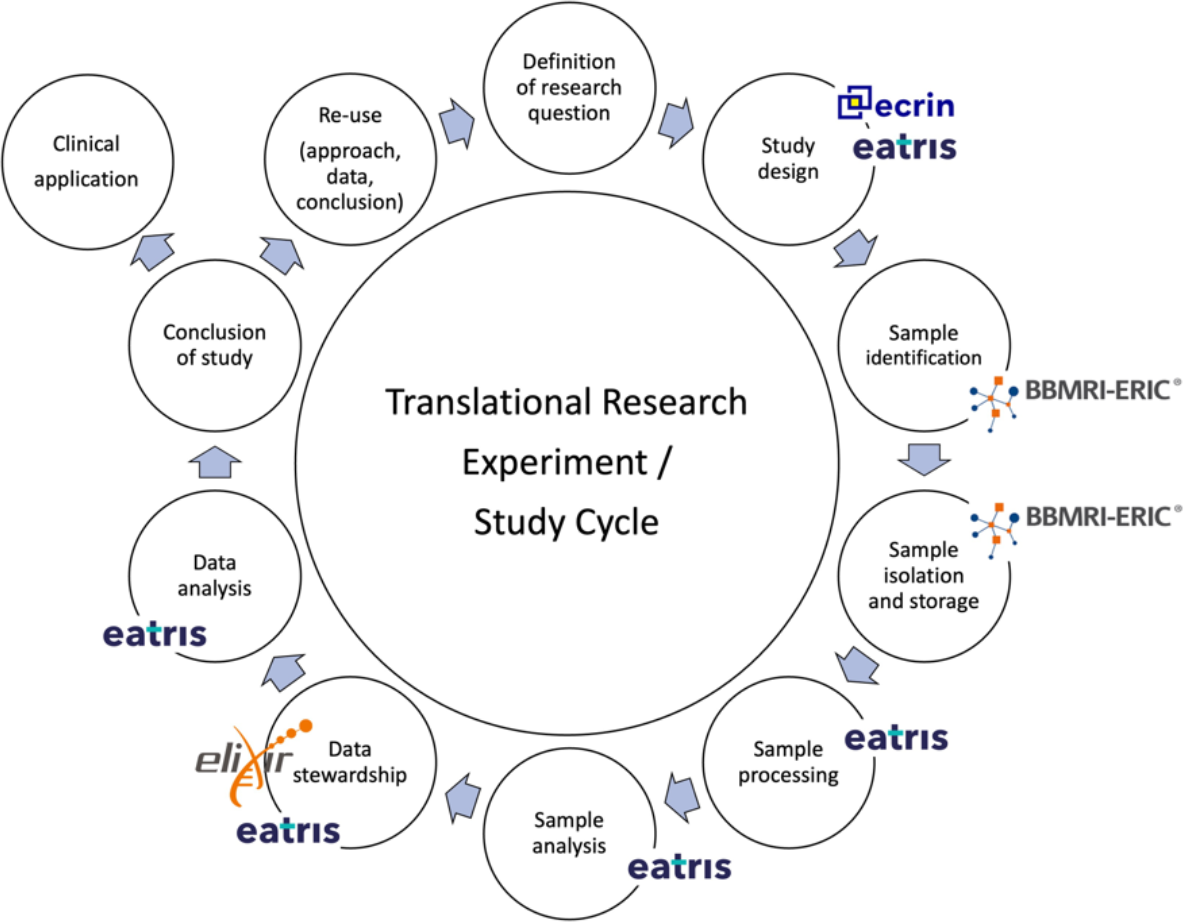
The translational biomedical process and involvement of selected biomedical Research Infrastructures as stakeholders in quality assessment. EATRIS is involved in quality assessment checkpoints along the translational trajectory. ECRIN, https://ecrin.org; BBMRI-ERIC, https://bbmri-eric.eu; Elxir, https://elixir-europe.org

The EATRIS-Plus project which was launched to deliver innovative scientific tools to support the long-term sustainability strategy of EATRIS as one of Europe’s key research infrastructures for Personalised Medicine. As part of the EATRIS-Plus project, a multi-omics dataset from about 100 healthy individuals was generated and compiled. The data of this pilot project are being made available for further analyses by stakeholders involved in biomedical research, healthcare, and drug development. The multi-omics study was based on samples collected from volunteers within the Czech Genome Project (https://czechgenome.iabio.eu/). Here we outline and discuss the efforts that the EATRIS-Plus initiative took to address comparative quality assessment (QA) of omics data generation and analysis. Procedures and results of this initiative have been compiled in the Multi-Omics Toolbox (MOTBX) (https://motbx.eatris.eu), (see below).

### Proficiency testing for assessing pre-analytical sample processing methods

Proficiency Testing (PT) programs are external quality assessment (EQA) tools that aim to evaluate the performance of laboratories in conducting specific measurements or tests, ensuring ongoing quality monitoring, and promoting the standardization of procedures to achieve greater consistency and reproducibility in results (3).

Participation in PT programs enables laboratories to gauge the strengths and weaknesses of their procedures by comparing performance and reproducibility of their results with those of their peers and allowing to validate and enhance performance, identify potential issues in testing and processing or technical problems with equipment or reagents, compare and harmonize methods and procedures, assess precision and accuracy, evaluate operator capabilities, provide staff education, and instill confidence in laboratory staff and users (4). PT programs can also offer valuable insights into method-trueness, particularly in cases where reference materials are not available, thus supporting analytical method validation (5).

PT programs play a crucial role in the quality management system of laboratories by ensuring consistent delivery of high-quality data and monitoring data reliability. Laboratories in which the results deviate significantly for the expected values (high z-scores) necessitate prompt corrective actions to be taken. To achieve this, implementing a comprehensive set of measures is essential. These measures encompass the utilization of standard operating procedures (SOPs) and validated protocols, incorporating internal quality controls such as reference materials and control charts, active participation in PT programs, and obtaining certification or accreditation according to recognized standards such as ISO15189/IEC, ISO/IEC 17025 and ISO 9001 (6). Indeed, it has been demonstrated that laboratories engaged in PT programs exhibit robust internal quality control procedures and consistently achieve better z-scores, (7, 8).

In the context of EATRIS-Plus, a PT program for biospecimen processing has been established by the Integrated BioBank of Luxembourg (IBBL, https://www.lih.lu/en/translational-medicine/translational-medicine-operations-hub/integrated-biobank-of-luxembourg-ibbl/; https://biospecimenpt.ibbl.lu/) to ensure the fitness-for-purpose of samples for the down-stream omics analysis. From 2020 to 2022 four EATRIS-Plus partners involved in omics analyses were enrolled in 26 processing schemes. The aim was to compare the efficiency of the processing methods with those of other laboratories, from different sectors worldwide, ensure the validity of the result, monitor, and improve performance by identifying potential problems, prove consistency of performance over time. **Figure 2** shows the case of an exemplary EATRIS facility with sequential participation in PT programs. In case specific problems are pinpointed in any of the determined parameters, the participating laboratory can take corrective measures, and this can lead to significant improvement in the quality of the sample processing. Participation in PT schemes represents a critical step also for omics analyses to ensure accurate, reliable, and trustworthy data.

**Figure 2.**
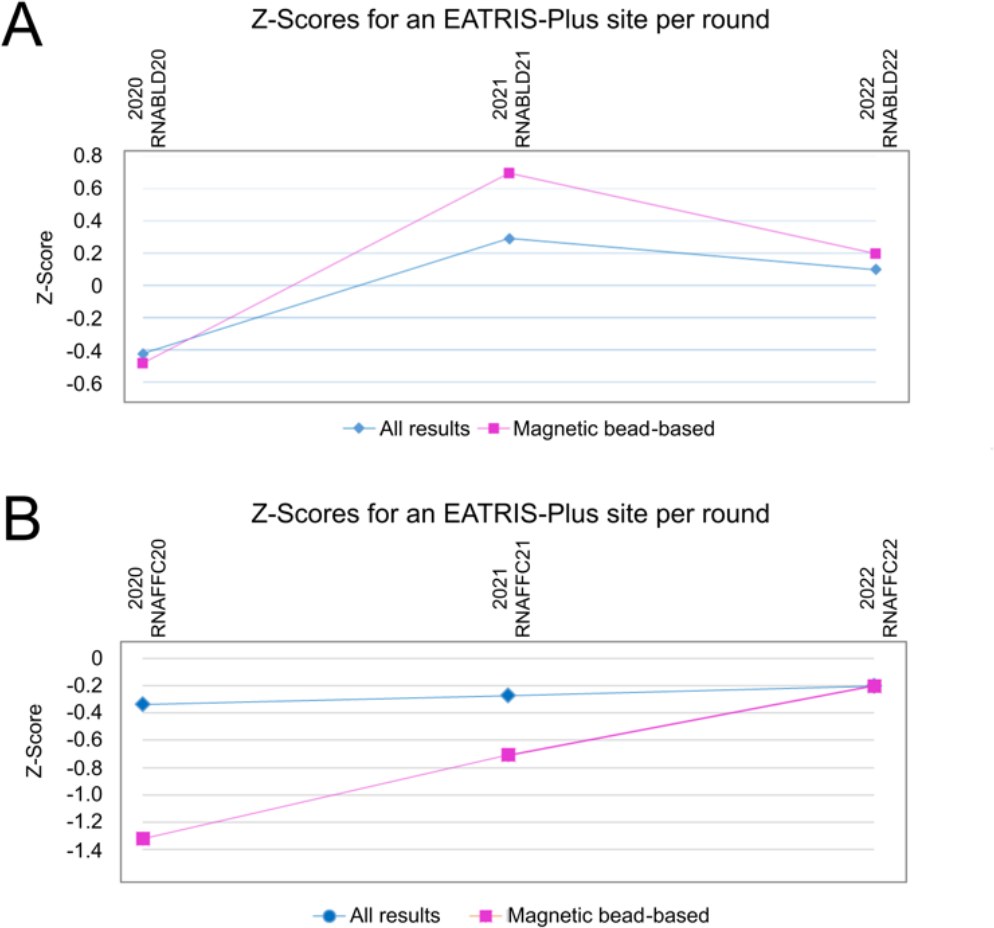
External quality assessment. History of z-Scores of one of the four EATRIS-Plus omics sites participating in the “RNA Extraction from Whole Blood” scheme” (A), and “RNA Extraction from FFPE Tissue” (B) scheme of the IBBL PT program from 2020 until 2022. In the long run, a large proportion of results giving rise to |z|>2 (more than 5%) and |z|>3 (more than 0.3%) indicates either a biased mean, or a standard deviation of the participant which is higher than the Proficiency Testing Standard Deviation. The participating site used a magnetic bead-based RNA isolation method, whereas the comparative score “All results” is an average z-Scores from all participants using magnetic bead-based, silica membrane-based or other RNA isolation methods.

### Multi-omics reference materials for quality assessment: commercial and research materials

The options for analysis of biological samples with omics platforms are numerous, and there is no undisputed or unchallenged “perfect” way. Largely depending on the research or clinical question and the available material, extraction methods may vary, and many technical and statistical approaches may be employed, sometimes leading to tremendous discrepancies in the results (e.g., (9)).

All facilities of the EATRIS-Plus project utilize high-quality reference material in the daily work routine for their respective omics analysis platforms to monitor changes of quality over time and from experiment to experiment (**Table 1**). Gradual deterioration or improvement of the data quality from the reference material as well as batch effects and other “unwanted noise” indicates that corrective measures are needed. In addition to these intra-lab QA, inter-laboratory QA can also be conducted. However, unless all labs use the exact same methodology (an ideal condition for direct lab-to-lab comparison), the comparative analysis usually relies on the identification of consensus features (e.g., concentration of analytes in a biofluid, detection of small variants in a DNA or specific transcripts in an RNA). In the case of multi-omics datasets management, batch detection and correction methods, and best practices for mitigation of unwanted noise are currently being developed (10, 11).

**Table 1.**
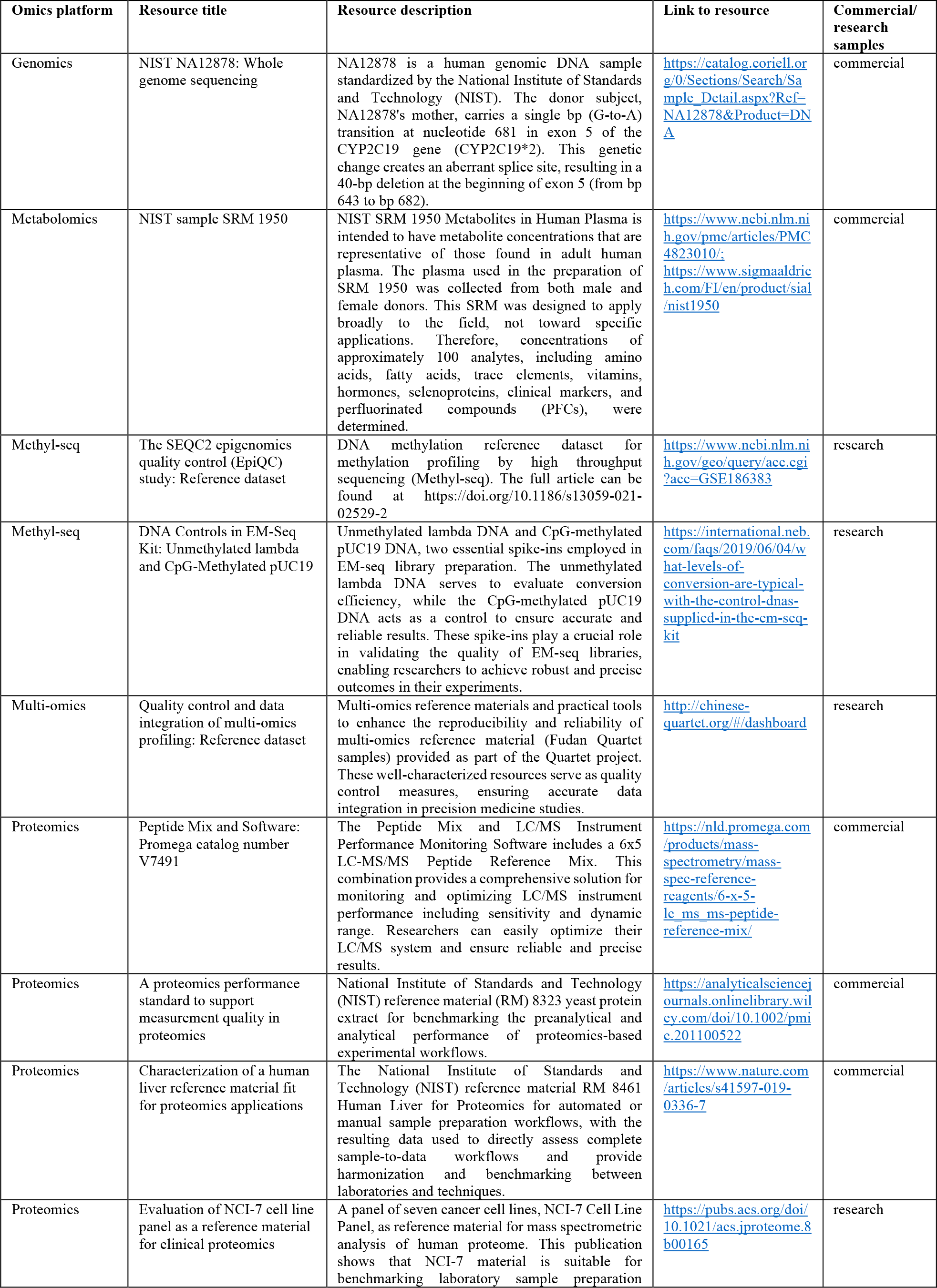
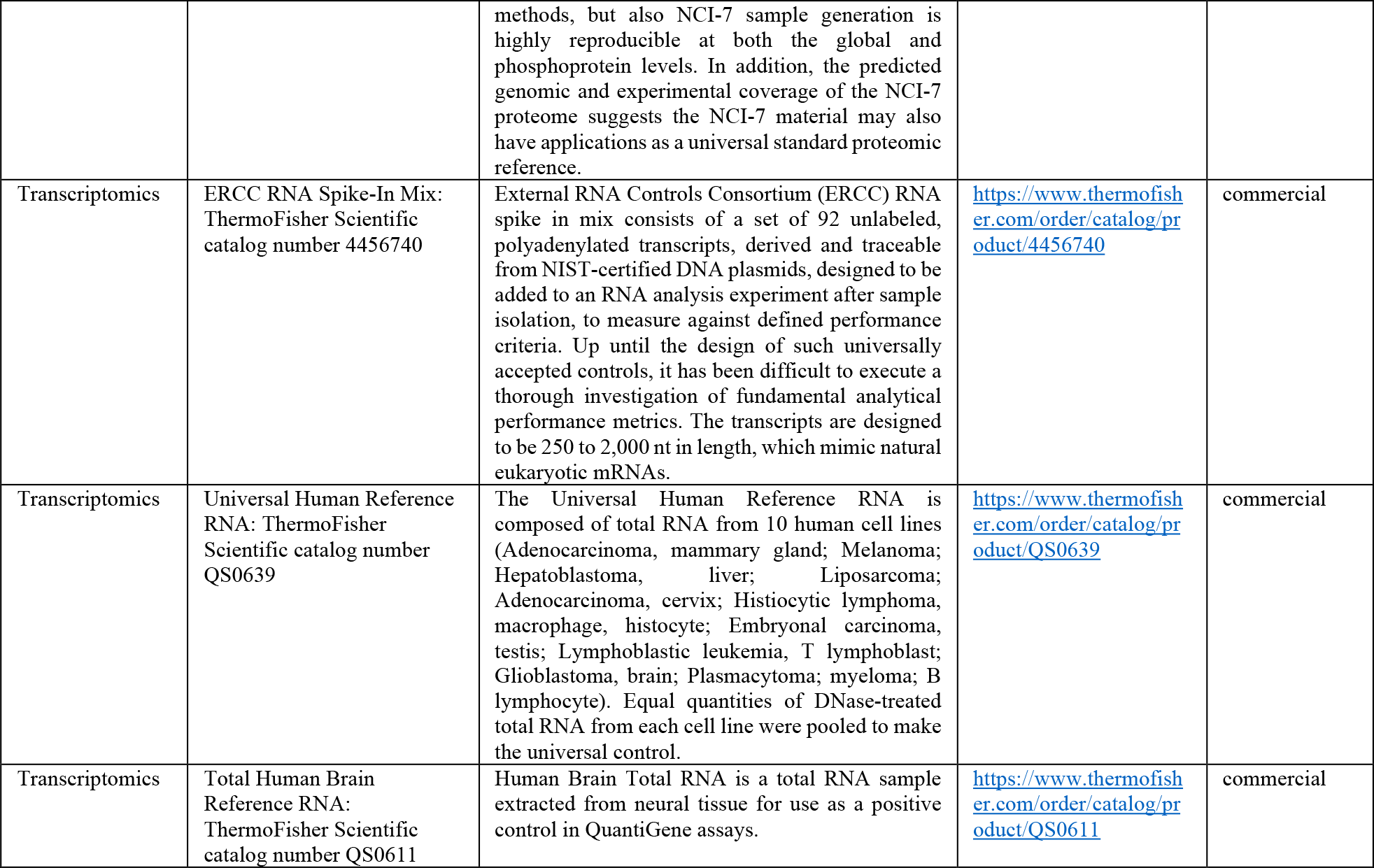
Reference material for omics technologies. Adapted from MOTBX website (https://motbx.eatris.eu).

While single omics reference materials are available and widely used as “ground truth” for the evaluation of the performance of the technologies and technology benchmarking (e.g. genomic DNA (12, 13), tumor-normal paired DNA (14-16), RNA (17), protein (18, 19) and metabolite reference material (20), multi-omics demands matched reference resources spanning DNA, RNA, proteins and metabolites. EATRIS Plus members have used both commercial and research reference materials during the project.

### Commercial omics reference material

To evaluate the process from acquisition of pre-extracted reference multi-omics material (RNA, DNA, metabolites, and proteins) to data generation, the EATRIS-Plus project employs both commercially available reference materials and research reference materials, namely the samples of the Fudan Quartet project (**Table 1**). Commercial quality references are regularly used by EATRIS Plus sites, usually for longitudinal quality assessment. These reagents are processed according to the procedures used at each site and allow for a comparison of data quality between laboratories. However, for certain omics technologies commercial reference materials are available to check only specific technical steps but not the whole process (from sample preparation to data analysis). For example, no reference materials specifically designed for differential protein expression analysis is commercially available, and usually in-house generated standards obtained my mixing at defined ratios commercially available individual tryptic digests are used as a proxy.

### Research omics reference material: The Fudan Quartet Project

In a PT effort aimed to assess the quality of the omics technology platforms of EATRIS-Plus partners, the EATRIS-Plus project acquired Quartet multi-omics reference materials from Fudan University (Shanghai, China). The Quartet reference materials encompass DNA, RNA, protein, and metabolites, all derived from B-lymphoblastoid cell lines obtained from a familial quartet consisting of parents and monozygotic twin daughters (21-24). The Quartet Project is a pivotal resource, providing “multi-omics ground truth”, best practices, and computational methods for objective assessment of proficiency and reliability of data generation processes in participating laboratories (**Figure 3**)

**Figure 3:**
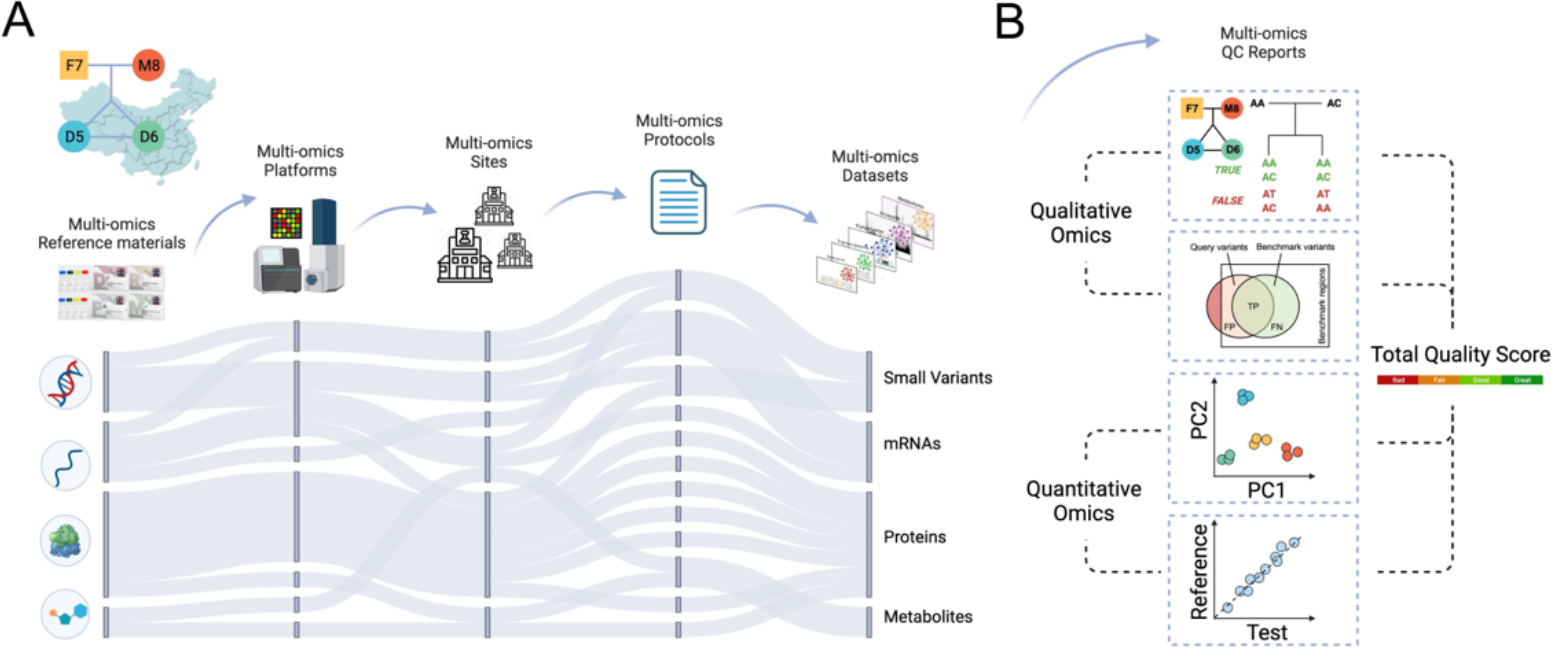
The Chinese Quartet reference materials and the quality assessment system. A, data generation by using Quartet reference materials across multiple platforms, sites and protocols. B, visual representation of several quality assessment parameters. The quality assessment system is embedded on the Quartet Data Portal (https://chinese-quartet.org/#/dashboard). A total quality score is calculated from several individual aspects of quality, for qualitative omics including Mendelian concordance rate and F1 score and for quantitative omics including signal-to-noise ratio (SNR) and relative correlation with reference dataset (RC) (23).

Quartet reference materials were dispatched to an EATRIS Plus coordinator site of this activity, aliquoted and then distributed to the various EATRIS laboratories for processing and analysis. Raw data resulting from these analyses were uploaded onto the Quartet Data Portal (https://chinese-quartet.org), where automated data analysis and reporting were conducted using publicly available workflows. EATRIS-Plus laboratories provided data for proteomics, metabolomics, DNAseq and RNAseq (**Figure 4)**. Three EATRIS sites performed also microRNAseq, which was manually analyzed by the Fudan Quartet team, as this workflow was not implemented in the Quartet Data Portal at the time this study was performed. Finally, one EATRIS Plus site performed and evaluated microRNA qRT-PCR for 170 microRNAs (data not shown).

**Figure 4.**
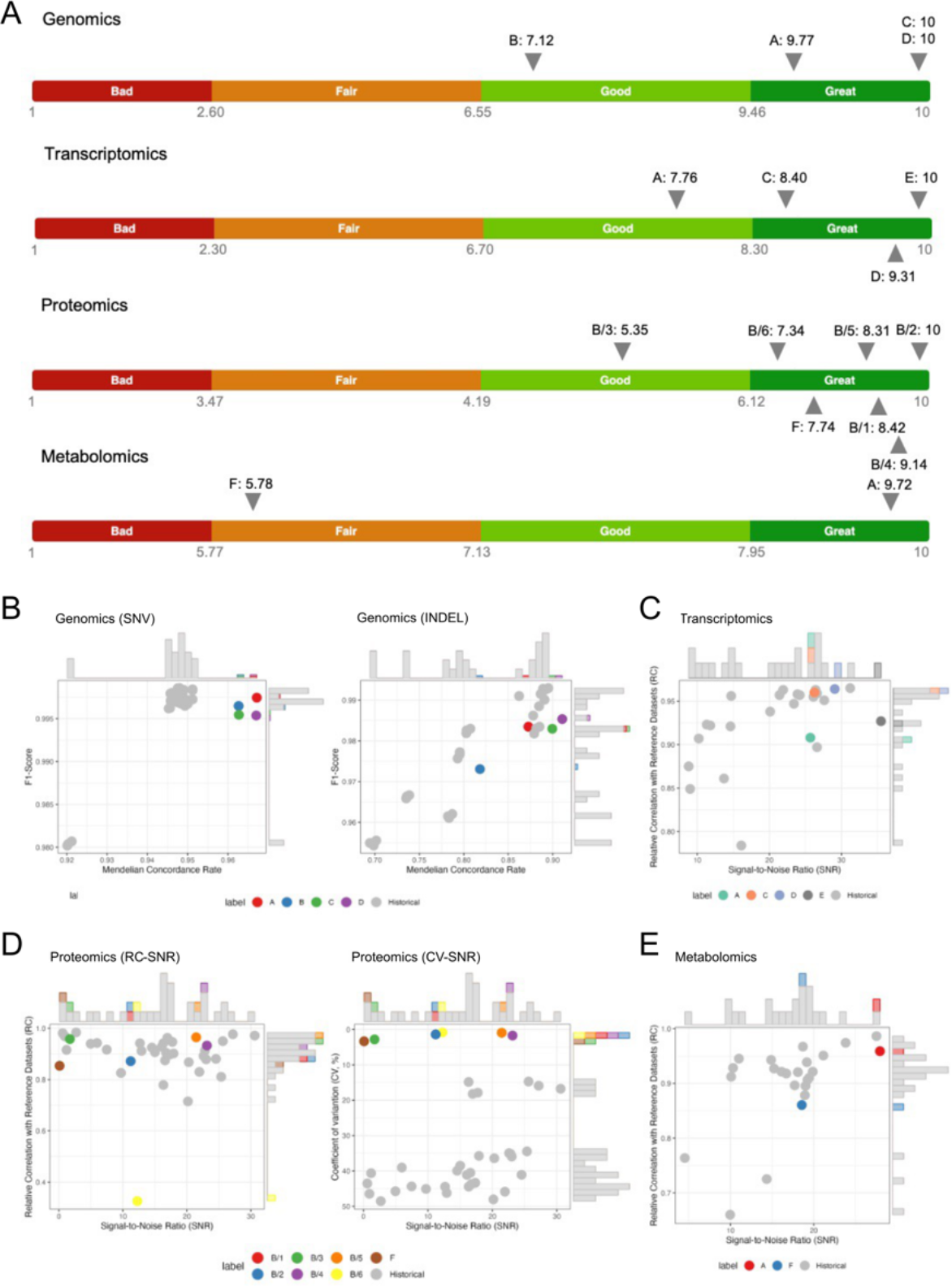
Quality assessment of multi-omics, multi-site, and multi-protocol datasets in proficiency testing. Chinese Quartet reference material was subjected to omics analysis in different EATRIS Plus facilities. A, the total quality scores of all datasets with ranking labels among all historical datasets in genomics, transcriptomics, proteomics and metabolomics. The label “Bad”, “Fair”, “Good” or “Great” manifests as the dataset ranking below the lower 20%, the 50%, the upper 20%, or above the upper 20% quantiles of the historical datasets. B-E, scatter plots of quality assessment results in genomics (B), transcriptomics (C), proteomics (D) and metabolomics (E) data; each datapoint shows the values of specific QC metrics across the samples in a given dataset. All historical datasets were colored as grey to be distinguished from the tested datasets. All scatter plots were added with frequency distribution bars to the marginal. Supplementary **Table S1** provides information on key aspects of the workflows. One potential use of the Fudan Quartet reference samples is demonstrated here for LC-MS proteomics analysis. Laboratory B evaluated six different LC-MS proteomics protocols, being different in fractionation column (performance (B2/4/6) versus endurance (B1/3/5)), MS mode (DDA (B1/2/3) versus DIA (B4/5/6)) and data analysis method (MSFragger (B1/2) versus PaSER (B3/4/5/6) (25-29). The quality of the proteomics output of these six protocols was assessed using the Fudan quality scoring yielding informative ranking that was subsequently used by the laboratory in their fit-for-purpose selection of workflows.

### Proficiency testing of EATRIS Plus facilities

The Quartet design provides both reference dataset-dependent and -independent QC metrics for quality assessment of multi-omics profiling (**Figure 4)**. For qualitative omics, the F1-score is a commonly used measure that takes into account both false positives and false negatives by computing a harmonic mean of precision and recall. Quality metrics for assessing reliability of DNA-seq, RNA-seq, proteomics, and metabolomics in terms of intra-batch proficiency and cross-batch reproducibility are assessed using ratio-based reference datasets (23). The Pearson correlation coefficient between the ratio-based expression levels of test datasets and reference datasets are used to describe the accuracy of quantitation. Signal-to-Noise Ratio (SNR) is used to investigate the differences between “cases” and controls. Overall, the platform allows to evaluate the quality of data coming from different participants, platforms, protocols, and analytical tools.

### Prerequisites for multi-omics analysis and integration

Implementing FAIR practices (1) at the individual omics level is essential to derive reproducible results from multi-omics data analyses. Most importantly, samples and data should be described by rich metadata (provenance). As acquired data and subsequent multi-omics analysis can be affected by technical factors related to sampling, processing, sample storage conditions (temperature, duration, thawing/freezing cycles), and measurement conditions (protocols, measurement order, possible measurement batches) documenting these factors can be key to reproducibility of research results. Additionally, intra- and inter-batch QC samples can help identify and adjust for batch effects (30). Finally, practices implementing FAIR Principles for Research Software can increase reproducibility of integrative multi-omics analyses (31-33).

The available analysis methods for multi-integration can be limited by type and completeness of data. While some integrative multi-omics methods can handle missing observations in some data modalities (30, 34), many methods require complete observations (35). Obtaining complete data for vertical integration is more challenging than in single omics experiments. Although methods that allow imputation of a low number of missing values are available (and the choice of imputation method should be guided by the mechanism that causes missing values, e.g. missing at random vs. not at random (36)), in the presence of numerous missing values, imputation is not advised.

### The EATRIS Multi-Omics Toolbox (MOTBX)

The multi-omics research community still faces a number of challenges impacting the biomarker development and implementation in clinical practice that need to be overcome: (a) poor levels of technological, analytical and data processing harmonization resulting in poor reproducibility, (b) poor data stewardship and compliance to the FAIR principles (1), (c) lack of understanding of the relationship between biomarkers belonging to different biological layers (transcriptomic, proteomic, metabolomic, epigenomic), (d) lack of reliable control reference values for these biomarkers, and (e) poor understanding of the actual clinical needs, resulting in limited clinical adoption (37).

In addition, information on omics reference material and multi-omics data integrative analysis and interpretation is fragmented in the knowledge space (38), and publicly available multi-omics profiling data are scarce.

Tackling these issues in a systematic way was one of the main objectives of the EATRIS-Plus project. This resulted in the development of the Multi-Omics Toolbox, MOTBX (https://motbx.eatris.eu).

MOTBX is a web-based open platform that is aimed to provide researchers, health professionals and other users with relevant information on quality standards and multi-omics resources to support their efforts towards the generation of high-quality data. The MOTBX core is structured into three sections: Omics Technologies, Quality Assessment, and Data, including analysis and FAIRification pipelines and tools (**Figure 5**).

**Figure 5.**
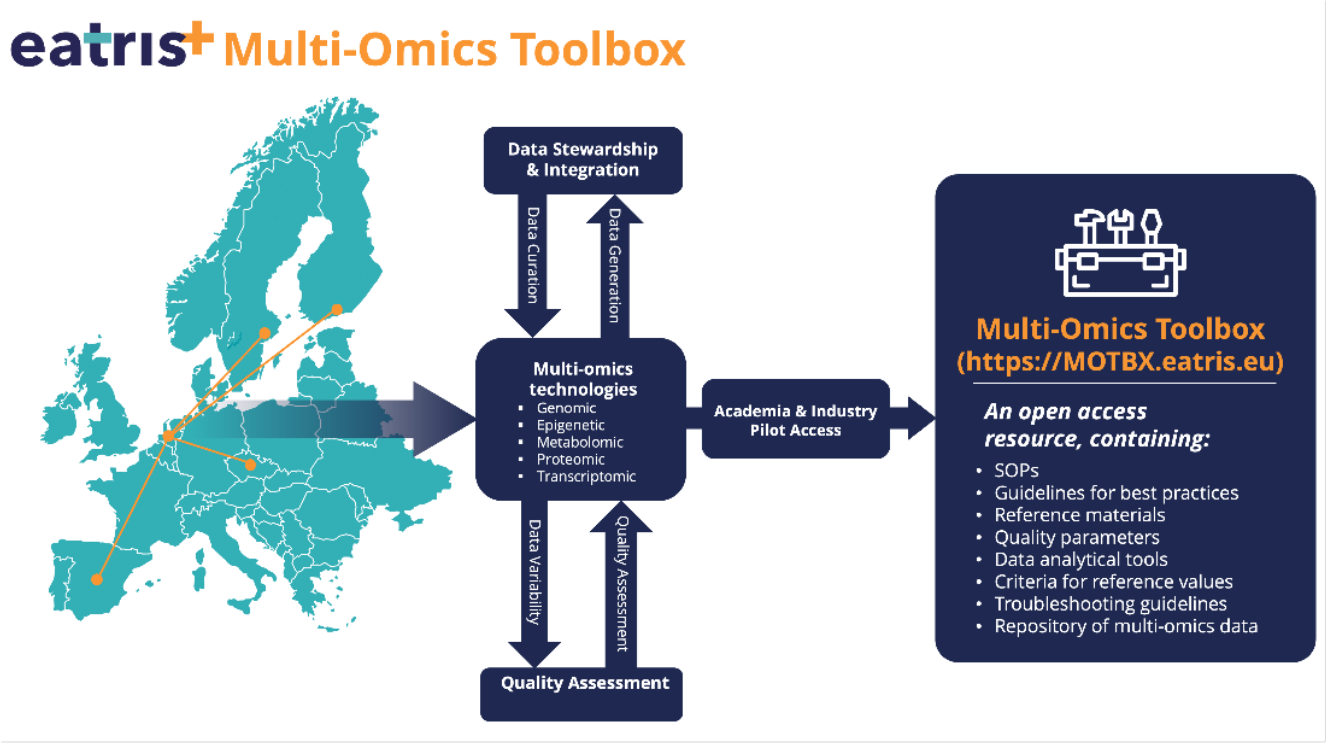
The Multi-Omics Toolbox. The Multi-Omics Toolbox (MOTBX, https://motbx.eatris.eu) is an open access knowledge hub for translational researchers supporting development, implementation and adoption of multi-omics approaches for personalized medicine, including quality assessment aspects. The MOTBX has been developed by EATRIS-Plus partners across Europe to support the translational biomedical research communities, as the result of collaborative work on data stewardship, the generation of a multi-omics dataset of healthy individuals, quality assessment studies, and the help and input of stakeholders from industry and academia.

MOTBX is foreseen to serve the scientific community as “meeting point” and a useful resource to obtain centralized access and information on multi-omics data generation, quality assessment and analysis, and FAIR principles. The user can access necessary elements required for performing high-quality research in the field of personalized medicine based on multi-modal biomarkers, such standard operating procedures (SOPs), clinical trial design, protocols, guidelines, reference materials and analytical tools.

In addition, the MOTBX includes education and training material, links to external resources, and a demonstrator multi-omics dataset, from an established Czech cohort of >100 healthy individuals, consisting of genome, epigenome, RNA and microRNA sequencing data, proteomics, and metabolomics data.

## Discussion

Specific and sensitive detection of biomarkers is one of the main pillars of personalized medicine. In this context, the procedures for sample harvesting and processing, data integration with phenotypic traits, analysis, and interpretation of results play a key role in the final decision on action. Depending on the outcome, this linear path may become circular or branch, when it is decided that further samples are needed, and other investigations are required to get a deeper understanding of the subject matter. The process requires quality assessment and quality control measures at every step: technology platforms should be validated, data collection and management transparent, data format and entries harmonized and electronically accessible, and the entire process including the analysis process and decision making should be thoroughly documented. European Biomedical Research Infrastructures have been developed since the early second decade of the 2000s to play a leading role in developing, applying, and disseminating quality assurance guidelines in different areas along the translational trajectory. For instance, BBMRI (www.bbmri-eric.eu) is addressing quality in human biological samples and sample storage; EuroBioimaging (www.eurobioimaging.eu) is providing image analysis solutions and imaging quality assessment guidelines; ELIXIR (www.elixir-europe.org) has been founded to develop and provide guidance for management and quality of life science data. In this context, EATRIS (www.eatris.eu) focuses on sample processing and analysis for personalized medicine applications.

In this work we share some of the initiatives EATRIS sites have participated in within the EATRIS Plus project. We show that proficiency testing of sample processing technologies and laboratory performance can and should be done at different steps of the translational process, as quality increases the chance of developing meaningful molecular tools. Longitudinal proficiency testing can lead to an improvement of laboratory performance, as it provides continuous feedback to the researcher on the quality of the sample processing. Application of commercial or research reference material serves multiple purposes, from assessing the quality of the data generation process at a given timepoint, the development of the quality over time, and the possibility to merge batches of data into larger datasets by applying statistical measures.

To demonstrate the applicability of the quality assessment strategies in the field of multi-omics applications mentioned above, EATRIS Plus generated a multi-omics dataset, consisting of DNA-seq, methyl-seq, RNA-seq, micro-RNAseq, microRNA qRT-PCR, proteomics, and metabolomics data of from about 100 healthy individuals (Czech cohort). We applied quality control and data integration strategies to create a holistic molecular representation of the individuals in the study. Together with references to literature on quality aspects of multi-omics data, links to analysis platforms and reference material, among others, these multi-omics data are available via MOTBX (https://motbx.eatris.eu), an open access multi-omics data platform that was created within the EATRIS Plus project. The small sample size used in this “demonstrator” does not allow us to define the obtained results as reference values for each omics in healthy population. However, EATRIS-Plus community aims to make the multi-omics dataset available for further growth by data integration with other datasets, thereby increasing its value for biomedical research and biomarker discovery. The MOTBX, as key public tool delivered by EATRIS-Plus community, is a live resource open to the entire research community that will be updated and implemented with new relevant content generated by qualified EATRIS partners, beyond the EATRIS-Plus project lifetime.

Since multi-omics approaches seek to discover interconnected signatures for sample classification and network identification between cross-omics features, QC metrics should be suitable for evaluating the performance of each omics type in terms of data generation and integration, and should be related to these two research objectives: (a) the integration of multi-omics information for more robust sample classifiers and, (b) the identification of multilayered interconnected molecular signatures are the major goals for multi-omics profiling. Quantitative benchmarks assessing these specific objectives, both within and between omics, are needed to drive advances in quality control and data integration (39). Thus, versatile, high-and accessible multi-omics quality reference materials paired with fit-for-purpose performance metrics are urgently needed (39-42). The use of multi-omics material is one of the measures that can be taken, but the definition of appropriate reference materials for omics implementation in clinical practice is one of the most critical aspects for omics adoption in health care systems for personalized medicine.

## Notes

### Competing Interest Statement

The authors have declared no competing interest.

### Summary of Updates

The author list was updated.

## References

1. Wilkinson MD, Dumontier M, Aalbersberg IJ, Appleton G, Axton M, Baak A, et al. The FAIR Guiding Principles for scientific data management and stewardship. Sci Data. 2016;3:160018.

2. van der Velde KJ, Singh G, Kaliyaperumal R, Liao X, de Ridder S, Rebers S, et al. FAIR Genomes metadata schema promoting Next Generation Sequencing data reuse in Dutch healthcare and research. Sci Data. 2022;9(1):169.

3. Miller WG, Jones GR, Horowitz GL, Weykamp C. Proficiency testing/external quality assessment: current challenges and future directions. Clin Chem. 2011;57(12):1670–80.

4. Brookman B, Stephenson N, Baumeister F, et al. Selection, Use and Interpretation of Proficiency Testing (PT) Schemes. Eurochem Guide. https://www.eurachem.org/images/stories/Guides/pdf/Eurachem_PT_Guide_2011.pdf 2011.

5. Analytical Method Committee TRSoC. The role of proficiency testing in method validation. Accreditation and Quality Assurance. 2010;15:73–9.

6. Meggendorfer M, Jobanputra V, Wrzeszczynski KO, Roepman P, de Bruijn E, Cuppen E, et al. Analytical demands to use whole-genome sequencing in precision oncology. Semin Cancer Biol. 2022;84:16–22.

7. Verderio P, Ciniselli CM, Gaignaux A, Pastori M, Saracino S, Kofanova O, et al. External Quality Assurance programs for processing methods provide evidence on impact of preanalytical variables. N Biotechnol. 2022;72:29–37.

8. Taverniers I, De Loose M, Van Bockstaele E. Trends in Quality in the Analytical Laboratory. II. Analytical Method Validation and Quality Assurance. Trends in Analytical Chemistry. 2004;23(8):532–52.

9. Shi L, Campbell G, Jones WD, Campagne F, Wen Z, Walker SJ, et al. The MicroArray Quality Control (MAQC)-II study of common practices for the development and validation of microarray-based predictive models. Nat Biotechnol. 2010;28(8):827–38.

10. Goh WWB, Wang W, Wong L. Why Batch Effects Matter in Omics Data, and How to Avoid Them. Trends Biotechnol. 2017;35(6):498–507.

11. Ugidos M, Tarazona S, Prats-Montalban JM, Ferrer A, Conesa A. MultiBaC: A strategy to remove batch effects between different omic data types. Stat Methods Med Res. 2020;29(10):2851–64.

12. Zook JM, Hansen NF, Olson ND, Chapman L, Mullikin JC, Xiao C, et al. A robust benchmark for detection of germline large deletions and insertions. Nat Biotechnol. 2020;38(11):1347–55.

13. Zook JM, McDaniel J, Olson ND, Wagner J, Parikh H, Heaton H, et al. An open resource for accurately benchmarking small variant and reference calls. Nature Biotechnology. 2019;37(5).

14. Jones W, Gong B, Novoradovskaya N, Li D, Kusko R, Richmond TA, et al. A verified genomic reference sample for assessing performance of cancer panels detecting small variants of low allele frequency. Genome Biology. 2021;22(1).

15. Deveson IW, Gong B, Lai K, LoCoco JS, Richmond TA, Schageman J, et al. Evaluating the analytical validity of circulating tumor DNA sequencing assays for precision oncology. Nature Biotechnology. 2021;39(9).

16. Fang LT, Zhu B, Zhao Y, Chen W, Yang Z, Kerrigan L, et al. Establishing community reference samples, data and call sets for benchmarking cancer mutation detection using whole-genome sequencing. Nat Biotechnol. 2021;39(9):1151–60.

17. Su Z, Łabaj PP, Li S, Thierry-Mieg J, Thierry-Mieg D, Shi W, et al. A comprehensive assessment of RNA-seq accuracy, reproducibility and information content by the Sequencing Quality Control Consortium. Nature Biotechnology. 2014;32(9).

18. Ivanov AR, Colangelo CM, Dufresne CP, Friedman DB, Lilley KS, Mechtler K, et al. Interlaboratory studies and initiatives developing standards for proteomics. Proteomics. 2013;13(6).

19. Friedman DB, Andacht TM, Bunger MK, Chien AS, Hawke DH, Krijgsveld J, et al. The ABRF Proteomics Research Group Studies: Educational exercises for qualitative and quantitative proteomic analyses. Proteomics. 2011;11(8).

20. Ulmer CZ, Ragland JM, Koelmel JP, Heckert A, Jones CM, Garrett TJ, et al. LipidQC: Method Validation Tool for Visual Comparison to SRM 1950 Using NIST Interlaboratory Comparison Exercise Lipid Consensus Mean Estimate Values. Analytical Chemistry. 2017;89(24).

21. Yu Y, Hou W, Liu Y, Wang H, Dong L, Mai Y, et al. Quartet RNA reference materials improve the quality of transcriptomic data through ratio-based profiling. Nat Biotechnol. 2023.

22. Yu Y, Zhang N, Mai Y, Ren L, Chen Q, Cao Z, et al. Correcting batch effects in large-scale multiomics studies using a reference-material-based ratio method. Genome Biol. 2023;24(1):201.

23. Zheng Y, Liu Y, Yang J, Dong L, Zhang R, Tian S, et al. Multi-omics data integration using ratio-based quantitative profiling with Quartet reference materials. Nat Biotechnol. 2023.

24. Yang J, Liu Y, Shang J, Chen Q, Chen Q, Ren L, et al. The Quartet Data Portal: integration of communitywide resources for multiomics quality control. bioRxiv.2022:2022.09.26.507202.

25. Meier F, Brunner AD, Frank M, Ha A, Bludau I, Voytik E, et al. diaPASEF: parallel accumulation-serial fragmentation combined with data-independent acquisition. Nat Methods. 2020;17(12):1229–36.

26. Meier F, Brunner AD, Koch S, Koch H, Lubeck M, Krause M, et al. Online Parallel Accumulation-Serial Fragmentation (PASEF) with a Novel Trapped Ion Mobility Mass Spectrometer. Mol Cell Proteomics. 2018;17(12):2534–45.

27. Yu F, Haynes SE, Teo GC, Avtonomov DM, Polasky DA, Nesvizhskii AI. Fast Quantitative Analysis of timsTOF PASEF Data with MSFragger and IonQuant. Mol Cell Proteomics. 2020;19(9):1575–85.

28. Xu T, Park SK, Venable JD, Wohlschlegel JA, Diedrich JK, Cociorva D, et al. ProLuCID: An improved SEQUEST-like algorithm with enhanced sensitivity and specificity. J Proteomics. 2015;129:16–24.

29. Demichev V, Messner CB, Vernardis SI, Lilley KS, Ralser M. DIA-NN: neural networks and interference correction enable deep proteome coverage in high throughput. Nat Methods. 2020;17(1):41–4.

30. Cuklina J, Lee CH, Williams EG, Sajic T, Collins BC, Rodriguez Martinez M, et al. Diagnostics and correction of batch effects in large-scale proteomic studies: a tutorial. Mol Syst Biol. 2021;17(8):e10240.

31. Barker M, Chue Hong NP, Katz DS, Lamprecht AL, Martinez-Ortiz C, Psomopoulos F, et al. Introducing the FAIR Principles for research software. Sci Data. 2022;9(1):622.

32. Chue Hong NP, Katz, D. S., Barker, M., Lamprecht, A-L, Martinez, C.,, Psomopoulos FE, Harrow, J., Castro, L. J., Gruenpeter, M., Martinez, P. A., Honeyman, T. FAIR Principles for Research Software version Zenodo; 2022.

33. de Visser C, Johansson LF, P. k, H. M P. N, Joeri van der Velde K, et al. Ten quick tips for building FAIR workflows. PLoS Comput Biol. 2023;19(9).

34. Argelaguet R, Arnol D, Bredikhin D, Deloro Y, Velten B, Marioni JC, et al. MOFA+: a statistical framework for comprehensive integration of multi-modal single-cell data. Genome Biol. 2020;21(1):111.

35. Flores JE, Claborne DM, Weller ZD, Webb-Robertson BM, Waters KM, Bramer LM. Missing data in multiomics integration: Recent advances through artificial intelligence. Front Artif Intell. 2023;6:1098308.

36. Wei R, Wang J, Su M, Jia E, Chen S, Chen T, et al. Missing Value Imputation Approach for Mass Spectrometry-based Metabolomics Data. Sci Rep. 2018;8(1):663.

37. Taube SE, Clark GM, Dancey JE, McShane LM, Sigman CC, Gutman SI. A perspective on challenges and issues in biomarker development and drug and biomarker codevelopment. J Natl Cancer Inst. 2009;101(21):1453–63.

38. Conesa A, Beck S. Making multi-omics data accessible to researchers. Sci Data. 2019;6(1):251.

39. Salit M, Woodcock J. MAQC and the era of genomic medicine. Nature Biotechnology2021.

40. Sené M, Gilmore I, Janssen JT. Metrology is key to reproducing results. Nature2017.

41. Wang X, Chambers MC, Vega-Montoto LJ, Bunk DM, Stein SE, Tabb DL. QC metrics from CPTAC raw LC-MS/MS data interpreted through multivariate statistics. Analytical Chemistry. 2014;86(5).

42. Beger RD, Dunn WB, Bandukwala A, Bethan B, Broadhurst D, Clish CB, et al. Towards quality assurance and quality control in untargeted metabolomics studies. Metabolomics. 2019;15(1).

